# D614G substitution enhances the stability of trimeric SARS-CoV-2 spike protein

**DOI:** 10.1101/2020.11.02.364273

**Authors:** Arangasamy Yazhini, Das Swayam Prakash Sidhanta, Narayanaswamy Srinivasan

## Abstract

SARS-CoV-2 spike protein with D614G substitution has become the dominant variant in the ongoing COVID-19 pandemic. Several studies to characterize the new virus expressing G614 variant show that it exhibits increased infectivity compared to the ancestral virus having D614 spike protein. Here, using *in-silico* mutagenesis and energy calculations, we analyzed inter-residue interaction energies and thermodynamic stability of the dominant (G614) and the ancestral (D614) variants of spike protein trimer in ‘closed’ and ‘partially open’ conformations. We find that the local interactions mediated by aspartate at the 614^th^ position are energetically frustrated and create unfavourable environment. Whereas, glycine at the same position confers energetically favourable environment and strengthens intra-as well as inter-protomer association. Such changes in the local interaction energies enhance the thermodynamic stability of the spike protein trimer as free energy difference (ΔΔG) upon glycine substitution is −2.6 kcal/mol for closed conformation and −2.0 kcal/mol for open conformation. Our results on the structural and energetic basis of enhanced stability hint that G614 may confer increased availability of functional form of spike protein trimer and consequent in higher infectivity than the D614 variant.

## Introduction

According to epidemiological surveillance of the disastrous COVID-19 pandemic, the causing agent *<u>s</u>*evere *<u>a</u>*cute *<u>r</u>*espiratory *<u>s</u>*yndrome coronavirus-2 (SARS-CoV-2) virus harbours mutations and associated with geographical-specific etiological effects (Brufsky, 2020; Mercatelli and Giorgi, 2020). Currently, three major variants of SARS-CoV-2 have been identified namely D614G in the spike protein, G251V in the non-structural protein 3 (NS3) and L84S in the ORF8 protein (Forster et al., 2020). In this article, we focused on the spike protein with D614G substitution. Spike protein of SARS-CoV-2 is a 1273aa long transmembrane glycoprotein and comprises of three modules namely a large ectodomain that protrudes from the surface, a single-pass transmembrane anchor and a short intracellular tail. The ectodomain has S1 and S2 regions responsible for host cell binding and viral-host membrane fusion, respectively. At the junction of S1 and S2 regions, S1/2 cleavage site is present and S2’ cleavage site is located in the S2 region. Depending on the orientation of a *<u>r</u>*eceptor *<u>b</u>*inding *<u>d</u>*omain (RBD) in the S1 region, protomers in the functional form of spike protein trimer adopts ‘open’ or ‘closed’ conformation (Cai et al., 2020; Walls et al., 2020; Wrapp et al., 2020).

Upon open conformation, RBD exposes ACE2 receptor binding regions and interacts with a peptidase domain of angiotensin-converting enzyme 2 (ACE2) receptor. This primary step clasps the virus on to the host surface (Lan et al., 2020; Yan et al., 2020). Studies on SARS-CoV have shown that subsequent proteolysis at S1/S2 cleavage site sheds S1 region from the spike protein and cleavage at S2’ site near fusion peptide causes a large conformational change in the S2 region. Such conformation change leads to an insertion of fusion peptide to host membrane and the formation of six-helix bundle. At this state, spike protein bridges viral envelope and host membrane. Hairpin-like bend in S2 region brings both membranes to close proximity for fusion following which genetic material gets released into the cytoplasm of the human cell (Belouzard et al., 2012). It is also noted that due to multi-basic nature of S1/2 cleavage site, the SARS-CoV-2 spike protein can be preactivated by furin enzyme during viral packaging (Shang et al., 2020). In contrast to SARS-CoV infection, this process reduces SARS-CoV-2 dependence on target cell proteases for the succeeding infection.

Therefore, mutations in the spike protein that influence the initial step for viral infection are associated with altered virus transmissibility and pathogenicity (Brufsky, 2020; Li et al., 2020). The ancestral spike protein with aspartate at 614^th^ position (S^D614^) has been asynchronously superseded by glycine substitution (S^G614^) world-wide. The dominant S^G614^ variant is shown to have higher infectivity than the S^D614^ variant (Korber et al., 2020). Concomitantly, other studies report that glycine substitution disrupts a salt bridge interaction between aspartate at 614^th^ position and lysine at 854^th^ position of an adjacent protomer and may contribute to higher frequency of open conformation than the S^D614^ variant (Cai et al., 2020; Yurkovetskiy et al., 2020; Zhang et al., 2020a). A recent cryo-EM study further reveals that the glycine substitution prevents premature shedding of S1 region (Zhang et al., 2020b). Along the similar line, our calculation of local interaction energies and free energy difference between aspartate and glycine variants of the spike protein, reported in this paper, suggests that glycine creates energetically favourable local environment. As a result, it strengthens the association of S1 and S2 regions of the same as well as adjacent protomer(s) and enhances overall stability of the spike protein trimer.

## Materials and Methods

### 3-D structural model for D614G variant of the spike protein trimer

We generated an *in silico* model for D614G variant of spike protein trimer using structure editing tool in UCSF chimera with default parameters (Pettersen et al., 2004). Side chains were optimized using SCWRL 4.0 program (Krivov et al., 2009). Two D614G variant models were generated corresponding to closed and partially open conformations of the spike protein trimer based on the reference cryo-EM structures available in the protein data bank (or PDB) entries 6VXX and 6VYB, respectively. It must be noted that although D614G variant structure is available (PDB code: 6XS6), we have not considered it in this analysis due to the absence of RBD domain in the solved structure.

### Calculation of frustration in the local interaction energy

Th effect of D614G substitution on local interaction energies was examined using Frustratometer algorithm (Parra et al., 2016). The underlying principle of the algorithm is that a native protein comprises of several conflicting contacts leading to local frustration. To examine frustrated contacts present in a protein, the algorithm systematically substitutes the residue type or alters chemical configuration of each interacting pair (including water-mediated interactions) and generates approximately 1000 structural decoys for a given contact (elaborated in Ferreiro et al. 2007). The extent of changes in the total interaction energy between native and structural decoys according to associative memory, water mediated, structure and energy model (AWSEM and implemented as a molecular dynamics algorithm, AWSEM-MD) decides whether the frustration of a concerned contact is minimal, neutral, or high. When the native energy is in the lower end of energy distribution of structural decoys, the contact is stabilizing and minimally frustrated (favourable) whereas when the native energy falls in between the energy distribution of structural decoys indicates the contact is neutrally frustrated. Native energy at higher end of the energy distribution of structural decoy indicates the contact is destabilizing and highly frustrated (unfavourable). Often, highly frustrated contacts signify functional constraints such as substrate binding, allosteric transitions, binding interfaces and conformational dynamics (Ferreiro et al., 2007, 2014).

Frustration of the contact is represented as frustration index, a Z-score of interaction energy of native contact with respect to the interaction energy distribution of structural decoys generated for that specific contact. Frustration index below −1 indicates the interacting pair is highly frustrated while the index between −1 to 0.78 or above 0.78 indicates the interacting pair is neutrally or minimally frustrated, respectively. Depending upon the nature of perturbation, frustration is referred as ‘mutational’, ‘configurational’ or ‘single-residue level’. In the mutational frustration, residue type is replaced by other residue types while in the configurational frustration, all possible interaction types between the native residue pairs were sampled through altering residue configuration. In case of single-residue level frustration, only a single residue is considered. The structural decoy set comprises of randomized residue type at that specific site and frustration index is calculated by evaluating changes in the protein energy upon altering the type of residue. In these three categories of frustration indices, only the concerned site/interaction is altered and the rest of the structure is maintained as native. In this study, we analyzed all categories of frustration indices for two variants of spike protein (S^D614^ and S^G614^) in the functional trimeric form.

### Calculation of thermodynamic stability of S^D614^ and S^G614^ variants

To study the effect of D614G variation on the thermodynamic stability of the spike protein trimer, we calculated free energy changes upon aspartate to glycine substitution using buildmodel function in FoldX (Schymkowitz et al., 2005). Five iterations of free energy calculations were carried out to obtain converged results (Tokuriki et al., 2007). Inferences of the results were derived from closed and partially open conformations of the spike protein trimer.

## Results and discussion

### D614G substitution increases the stability of spike protein trimer

As amino acid substitution alters local chemical environment, we probed D614G effect on the energetics of local inter-residue interactions. This can be quantified as local frustration of a residue or inter-residue interactions. We calculated frustration index of residues and interresidue interactions for two variants of spike protein *viz.* aspartate or glycine at the 614^th^ position. Result shows that frustration index of aspartate in the spike protein (S^D614^) is −1.25, – 1.25 and −1.30 for three protomers in the closed conformation (red lines in Figure 1A). The frustration index of aspartate in the partially open conformation is −1.24, −1.31 and −1.28 for three protomers (red lines in Figure 1B). Hence, in both the conformations aspartate is highly frustrated. Conversely, in glycine variant (S^G614^), the residue is neutrally frustrated with frustration index of −0.48, −0.42 and −0.46 for protomers in the closed conformation and −0.50, −0.35 and −0.37 for protomers in the partially open conformation (blue lines in Figure 1). This result implies that residue frustration at the 614^th^ position has become neutral upon glycine substitution.

**Figure 1.**
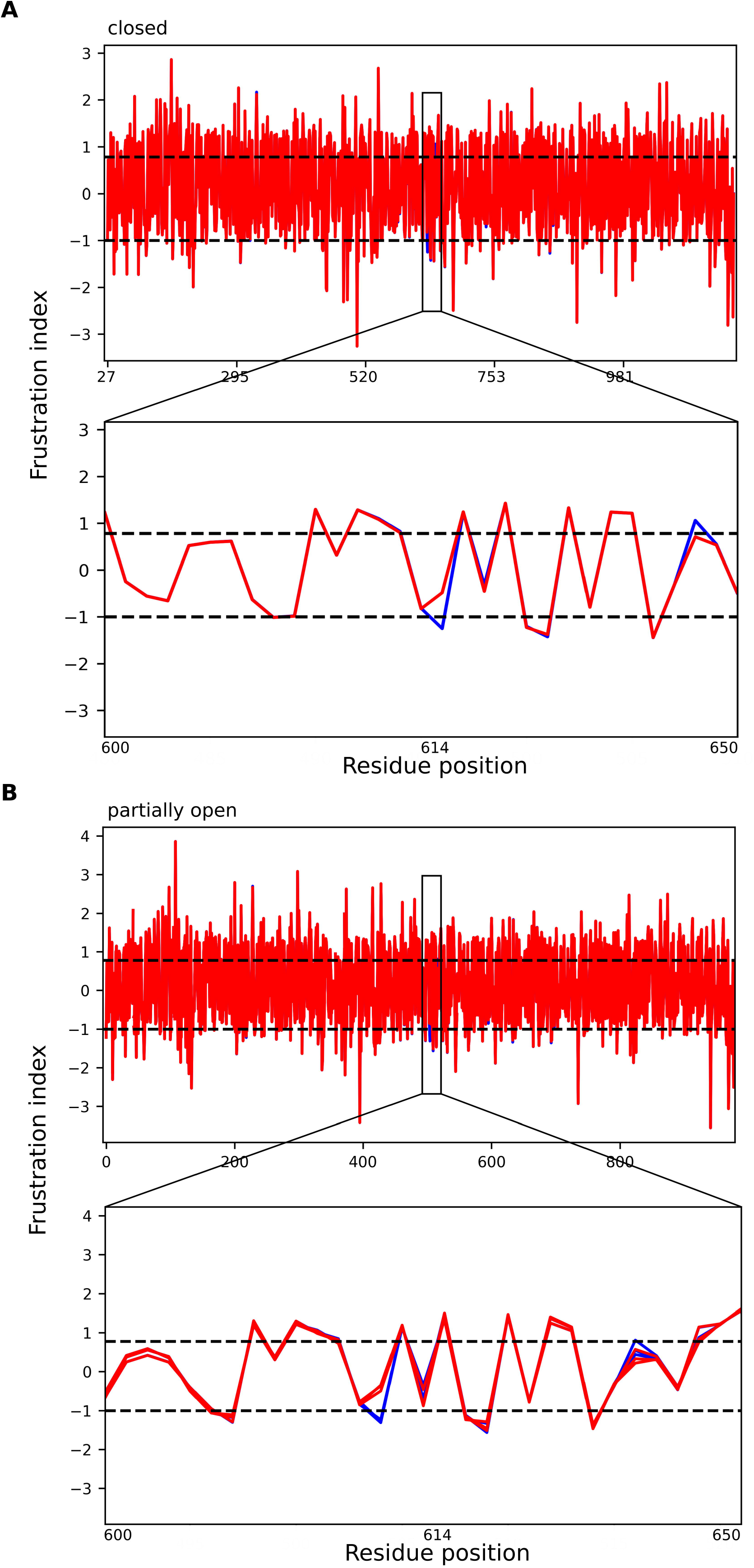
Frustration index of residues in S^D614^ and S^G614^ variants. Line plots show residue-wise frustration index of trimeric spike protein having aspartate (blue) or glycine (red) at 614^th^ position in closed (A) and partially open conformation (B). Horizontal dashed lines within the plots separate ranges with minimal (>0.78, top), neutral (−1 to 0.78, middle) and high (< −1, bottom) frustration index. In the bottom panel, region around 614^th^ position (600-650) is zoomed to indicate the loss of frustration in the local interaction energies upon aspartate to glycine substitution.

In the spike protein of both conformations (S^D614^), aspartate is involved in intra-protomer contact (with residues Ser591, Gly593 and Gly594) as well as in inter-protomer contact (with Asn616, Arg646, Ser735, Thr859 and Pro862) through direct, long-range electrostatic or water-mediated interactions. Mutational frustration index indicates that all the 8 contacts are highly frustrated (Figure 2, top panel). However, in the closed conformation of S^G614^ variant, glycine interacts with Phe318, Leu611 and Cys649 of the same protomer and Pro862 of the adjacent protomer. Except inter-protomer contact through Pro862, all three intra-protomer contacts are minimally frustrated (Figure 2, top left panel). Likewise, in the partially open conformation of S^G614^ variant, glycine has the same contact pattern as observed in the closed conformation along with one additional contact to Val860. Of these five contacts, three are minimally frustrated and two are highly frustrated (Figure 2, top right panel). Overall, the number of contacts as well as the number of highly frustrated contacts are reduced upon aspartate to glycine substitution. Notably, glycine forms a greater number of minimally frustrated contacts indicating it creates more favourable environment around the 614^th^ position compared to aspartate.

**Figure 2.**
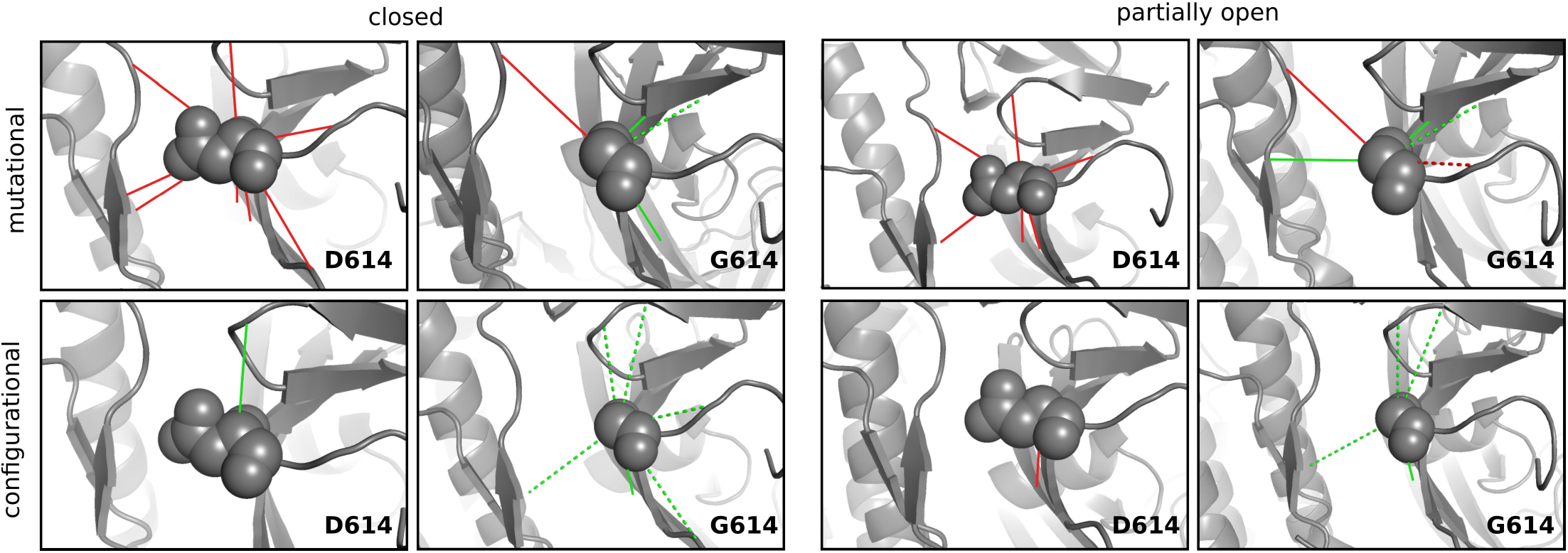
Frustration in the local interaction energies of S^D614^ and S^G614^ variants. Shown are the mutational (top panel) and configurational (bottom panel) frustrations exist in the inter-residue interactions formed by aspartate or glycine at 614^th^ position. Minimally and highly frustrated interactions are indicated by green and red lines, respectively. Water-mediated interactions are represented as dashed lines and the variant residue is shown as sphere. Results for closed and partially open conformation were shown for a protomer (chain ID: A) in the trimer and similar patterns are observed for the other two protomers (Supplementary Table S1).

Next, we calculated configurational frustration index that indicates how favourable the native contact between two residues relative to other possible contacts those two residues can have. Results show that in the closed conformation, aspartate (S^D614^) has one minimally frustrated contact with Arg646 (Figure 2, left bottom panel). Whereas, glycine (S^G614^) has six minimally frustrated contacts with residues Ser591, Gly593, Asn616, Thr645 and Arg646 of the same protomer and Thr859 of a preceding protomer in the clock-wise direction (Figure 2, left bottom panel). Similar trend is seen for partially open conformation in which aspartate (S^D614^) has a highly frustrated contact with Gly593 while glycine (S^G614^) has the same contacts but minimally frustrated besides three minimally frustrated contacts with other residues (Thr645, Arg646 and Thr859) (Figure 2, right bottom panel). These observations are common among three protomers present in the spike protein trimer (Supplementary Table S1). Hence, glycine has more favourable contacts than aspartate. Overall, calculations of single residue, mutational and configurational frustrations reveal that glycine substitution modified local interaction energy in the favourable direction.

If the reduction of frustration in the local interaction energies is significant upon aspartate to glycine substitution, it can have an influence on the thermodynamic stability of the spike protein trimer. To examine this, we calculated difference in the total free energy of trimer between S^D614^ and S^G614^ variants using FoldX package (Schymkowitz et al., 2005). Results show that the free energy difference (ΔΔG) is −2.6 kcal/mol for the closed conformation and – 2.0 kcal/mol for the partially open conformation. It suggests that the stabilizing effect of glycine substitution in the local environment markedly increases the overall stability of spike protein trimer. Together, these results imply that the enhanced stability of S^G614^ may confer increased availability of functional form of spike protein trimer and consequent in higher infectivity compared to the S^D614^ as observed in the recent experimental studies (Korber et al., 2020; Zhang et al., 2020a, 2020b).

## Conclusions

The increasing severity in public health and economic crisis builds urgency to develop therapeutic intervention against COVID-19 infection at the earliest. Currently, the dominance of D614G variant of SARS-CoV-2 spike protein that is being intensively studied across the globe for COVID-19 prophylaxis and treatment invites special attention. In this study, we demonstrate using *in-silico* approaches that glycine substitution at 614^th^ position changes the local environment from energetically frustrated into favourable for contacts present within as well as between protomer(s). Consequently, the free energy of S^G614^ is lower than that of S^D614^ and hence local changes in the interaction energies at the 614^th^ position in each protomer have a significant effect on the overall thermodynamic stability of the spike protein trimer. This finding bestows to our knowledge on the mechanism of increased transmissibility of S^G614^.

## Acknowledgements

This research is supported by Mathematical Biology program and FIST program sponsored by the Department of Science and Technology and also by the Department of Biotechnology, Government of India in the forms of IISc-DBT partnership programme. Support from UGC, India – Centre for Advanced Studies and Ministry of Human Resource Development, India is gratefully acknowledged. N.S. is a J. C. Bose National Fellow.

## Supplementary table legend

**Supplementary Table S1. The table contains details of frustration index of inter-residue contacts present at 614th position of spike protein trimer in closed and partially open conformations.** Table S1A in Sheet 1 provides mutational frustration index of contacts present at the 614th position in the ancestral (D614) and dominant (G614) variants of the spike protein trimer. In table S1B at sheet 2, configurational frustration index of those contacts in the ancestral (D614) and dominant (G614) variants has been provided. Frustration state namely minimally, neutral or highly represents that the contact is energetically favourable, neutral or unfavourable, respectively.

